# Template switching causes artificial junction formation and false identification of circular RNAs

**DOI:** 10.1101/259556

**Authors:** Chong Tang, Tian Yu, Yeming Xie, Zhuqing Wang, Hayden McSwiggin, Ying Zhang, Huili Zheng, Wei Yan

**Affiliations:** Department of Physiology and Cell Biology, University of Nevada, Reno School of Medicine, 1664 North Virginia Street, MS575, Reno, NV 89557; Department of Biology, University of Nevada, Reno, 1664 North Virginia Street, MS575, Reno, NV 89557, USA

**Keywords:** circular RNA, terminal transferase, RNA fusion, aging, experimental artifact, reproducibility

## Abstract

Hundreds of thousands of putative circular RNAs have been identified through deep sequencing and bioinformatic analyses. However, the circularity of these putative RNA circles has not been experimentally validated due to limited methodologies currently available. We reported here that the template-switching capability of commonly used reverse transcriptases (e.g., SuperScript II) leads to the formation of artificial junction sequences, and consequently misclassification of large linear RNAs as RNA circles. Use of reverse transcriptases without terminal transferase activity (e.g., MonsterScript) for cDNA synthesis is critical for the identification of physiological circular RNAs. We also report two methods, MonsterScript junction PCR and high-resolution melting curve analyses, which can reliably distinguish circular RNAs from their linear forms and thus, can be used to discover and validate true circular RNAs.

**Significance Statement:** The vast majority of circular RNAs were identified through computational detection of junction sequences in the deep sequencing reads because these unique fusion sequences represent back-splicing events. We found that artificial junction sequences could be formed through template switching (TS) when MMLV-derived reverse transcriptases, e.g., SuperScript II, are used to synthesize cDNAs. Thus, many of the reported circular RNAs may not be RNA circles, but rather experimental artifacts. Fake circular RNAs can be avoided by using reverse transcriptases without terminal transferase activity (e.g., MonsterScript) for cDNA synthesis. We developed two novel methods, MonsterScript junction PCR and high-resolution melting curve analyses, for distinguishing circular RNAs from their linear form.

---

Since their discovery ~two decades ago (1), thousands of circular RNAs have been reported in various species (2, 3). Two steps critical to circular RNA identification include RNase R treatment of total RNAs prior to cDNA synthesis, and bioinformatic identification of junction sequences indicative of self-ligation of mRNA fragments (2, 4, 5). It has been shown that RNase R treatment cannot remove all linear RNAs, especially those with complex secondary structures or chemical modifications (e.g., m6A) (6). Variations in RNases R activities among batches, manufactures, and digestion conditions, can also affect the efficiency of linear RNA removal. Algorithms used to computationally identify back-splicing patterns are mostly based on features of the junction sequences in circular RNAs reported in the literature although the circularity of the vast majority of the reported circular RNAs has not been validated experimentally. Northern blot, given its independence of RNase R treatment, has been regarded as a “gold standard” for detecting circular RNAs; however, the detection sensitivity is relatively low compared to PCR-based methods, and the specificity is also limited due to the fact that junction probes can still detect linear RNAs containing partial junction sequences. In our attempt to identify and validate true RNA circles, we discovered that the template-switching capability of commonly used reverse transcriptases could lead to false positive identification of junction sequences and consequently misclassification of large linear RNAs as RNA circles. We report here that the use of MonsterScript, a thermostable reverse transcriptase with neither RNase H nor terminal transferase activity, for reverse transcription during library construction and PCR-based analyses, can minimize false positive identification of circular RNAs, and the circularity of RNAs can be validated using both MonsterScript junction PCR and high-resolution melting curve analyses.

## Results and Discussion

### Template switching caused by terminal transferase activity of commonly used reserve transcriptases

In addition to DNA polymerase activity, the commonly used reverse transcriptases, including both MMLV- [e.g., SuperScript II (Thermo Fisher), ProtoScript II (NEB), PrimeScript (Clontech), GoScript (Promega), etc.] and AMV-derived ones, all possess terminal transferase activity, which can add several non-templated nucleotides to the 3’ end of cDNAs, leading to *template switching* (TS) during reverse transcription (RT) (**Fig. 1A**). The TS events can generate artificial junctions derived from the same linear RNA templates and these linear RNAs would be erroneously identified as RNA circles during bioinformatic annotation (**Fig. 1A**). We tested numerous reverse transcriptases and also modified the RT steps by utilizing Mung Bean nuclease, exonuclease VII or T7 endonuclease to trim the cDNA 3’ ends to minimize TS events, but all failed to distinguish between the circular and linear forms of the same control RNA *(SI Appendix, Fig. S1).* Therefore, use of MMLV-derived reserves transcriptases during RT inevitably generates artificial junction sequences as long as the linear RNAs are present in the reactions.

**Fig. 1.**
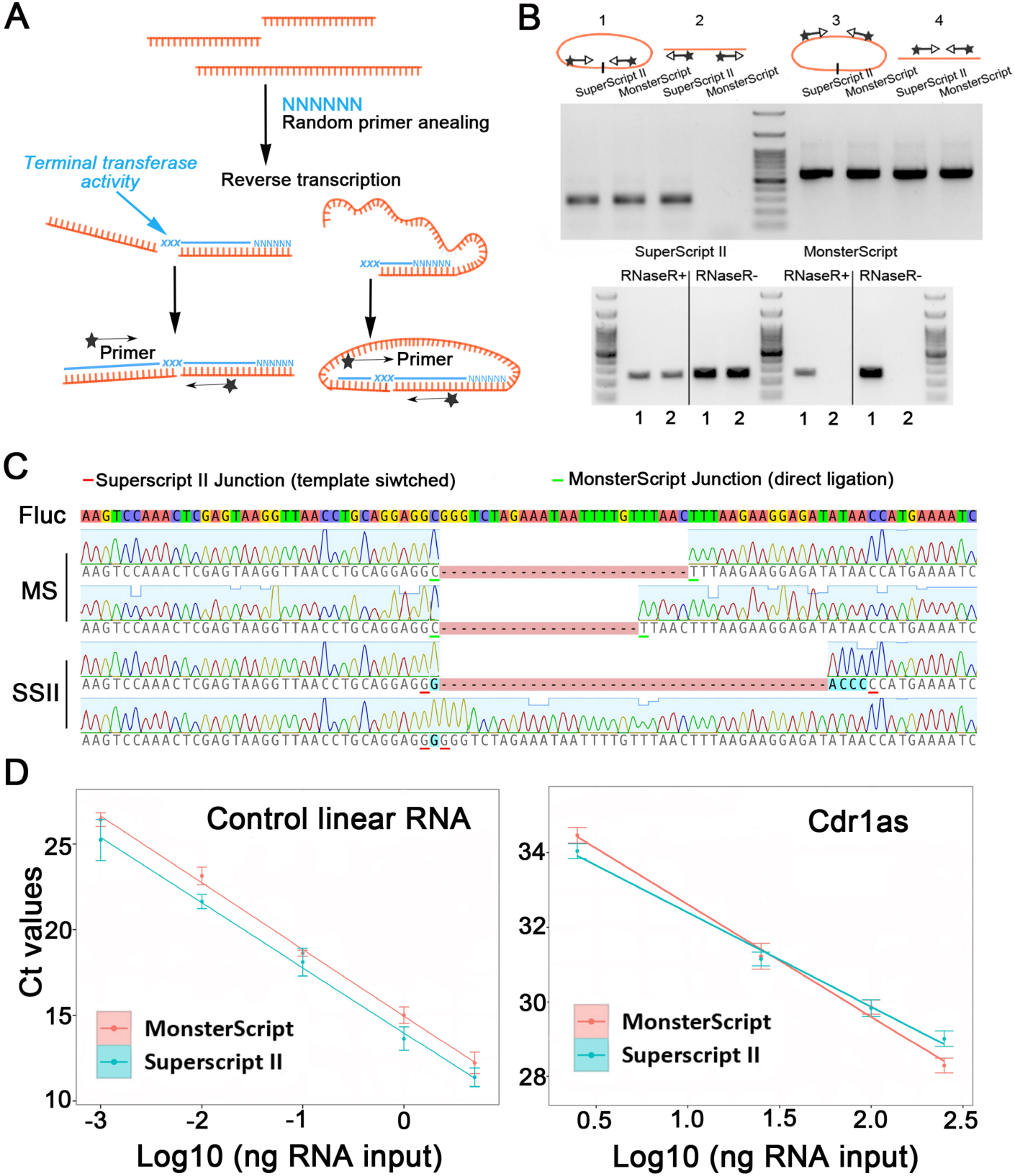
Junction formation due to template-switching (TS) when commonly used reverse transcriptases (e.g., SuperScript II) are used for cDNA synthesis. (**A**) Schematic presentation of the mechanism through which the terminal transferase activity of MMLV-derived reverse transcriptases (e.g., SuperScript II) can add several non-templated nucleotides to the 3’ end of cDNA, leading to TS during reverse transcription (RT). The TS events can generate artificial junctions derived from the same linear RNA templates and these linear RNAs would be erroneously identified as RNA circles in bioinformatic analyses. (**B**) PCR detection of the junction sequences from SuperScript II or MonsterScript RT products of both control circular (sample 1) and linear (sample 2) RNAs using convergent and divergent primers, respectively (upper left half panel). PCRs using internal primers were used as loading controls (upper right half panel). The lower panels show results after RNase R treatment of the control linear and circular RNAs. (**C**) Sanger sequencing of the PCR products. Note that the divergent PCR products of SuperScript II cDNAs from the control linear RNA contained insertions and mutations resulting from TS events, whereas both SuperScript II and MonsterScript convergent PCR products were from the true RNA ligation junctions of the control circular RNA. Fluc, control luciferase RNA; MS, MonsterScript; SSI I: SuperScript II. (**D**) Reverse transcriptase efficiency assays using the control linear RNA (left panel) and a previously validated circular RNA, *Cdr1as* (right panel). RNAs were reverse transcribed using SuperScript II and MonsterScript, followed by qPCR using the internal primers (for control linear RNA) and junction primers (for Cdr1as), respectively. CT values were plotted against log 10 values of RNA inputs and SuperScript II and MonsterScript displayed similar sensitivity in detecting either linear or circular RNAs.

### MonsterScript does not cause template switching during reverse transcription

Interestingly, we found that template switching did not occur when MonsterScript, a thermostable reverse transcriptase with neither RNase H nor terminal transferase activity, was used for reverse transcription. We first used MonsterScript to replace SuperScript II in 5’ RACE assays, and found that MonsterScript failed to detect the 5’ ends *(SI Appendix, Fig. S2*). Given that the 5’ RACE assay relies on the template switching property of SuperScript II to detect the 5’ end sequences, this result suggests that MonsterScript does not cause template switching. With control linear *(in vitro* transcription products of a luciferase plasmid) and circular RNAs (self-ligation products of linear luciferase RNA treated with RNase R) as templates, convergent primers successfully amplified the junction region of both SuperScript II and MonsterScript RT products of the control circular RNA (**Fig. 1B**). However, PCR using divergent primers consistently amplified the SuperScript II, but not the MonsterScript RT products of the control linear RNA (**Fig. 1B**). Sanger sequencing confirmed that the divergent PCR products of SuperScript II cDNAs from the control linear RNA contained insertions and mutations resulting from TS events (**Fig. 1C**).

RNase R has been routinely used to enrich circular RNAs. However, we noticed that RNase R treatment could not completely eliminate false positive detection of TS-derived junctions in divergent PCR on Superscript II RT products of control linear RNA despite our many attempts using RNase R from different vendors (**Fig. 1B**, **lower panels**). This is consistent with a recent report showing that RNase R treatment cannot remove all linear RNAs, especially those with complex secondary structures or chemical modifications (e.g., m6A)(6). In fact, RNase R treatment appeared to decrease the detection efficiency of MonsterScript junction PCR, probably due to hydrolysis of circular RNAs by combined catalytic activity of temperature and magnesium ions (7).

To see whether the difference reflects lowered detection sensitivity of MonsterScript, we performed junction PCR using SuperScript II and MonsterScript RT products from the control linear RNA and a previously validated circular RNA, *Cdr1as*(3). Similar robustness was observed between SuperScript II and MonsterScript (**Fig. 1D**), suggesting that the failure for MonsterScript to detect false positive junction sequences is not due to lower sensitivity compared to SuperScript II.

To further demonstrate the extent to which TS events occur during cDNA synthesis, we constructed libraries using SuperScript II and MonsterScript and performed RNA-Seq on the ERCC RNA Spike-In Mix (Invitrogen), which contained 96 synthetic linear RNAs with variable lengths, GC contents and amounts. Under similar sequencing depths (~45 million reads), both methods yielded similar read counts (**Fig. 2A**). Based on junction sequences detected, 20 artificial circular RNAs were identified from libraries constructed using SuperScript II, whereas none was found in the MonsterScript libraries (**Fig. 2B**). Certain sequencing depth was required to start to see noises (i.e., artificial circular RNAs) and the noise-to-signal ratios increased up to 10^65^ reads, and then started to decrease with the increasing sequencing depth in SuperScript II libraries of the 96 spike-in linear RNAs (**Fig. 2C**). In contrast, the noise-to-signal ratios were constantly low in MonsterScript libraries (**Fig. 2C**). These data suggest that MonsterScript, unlike the most commonly used MMLV-derived reverse transcriptases, e.g., SuperScript II, does not cause template switching and thus, can be used to identify biological rather than artificial circular RNAs.

**Fig. 2.**
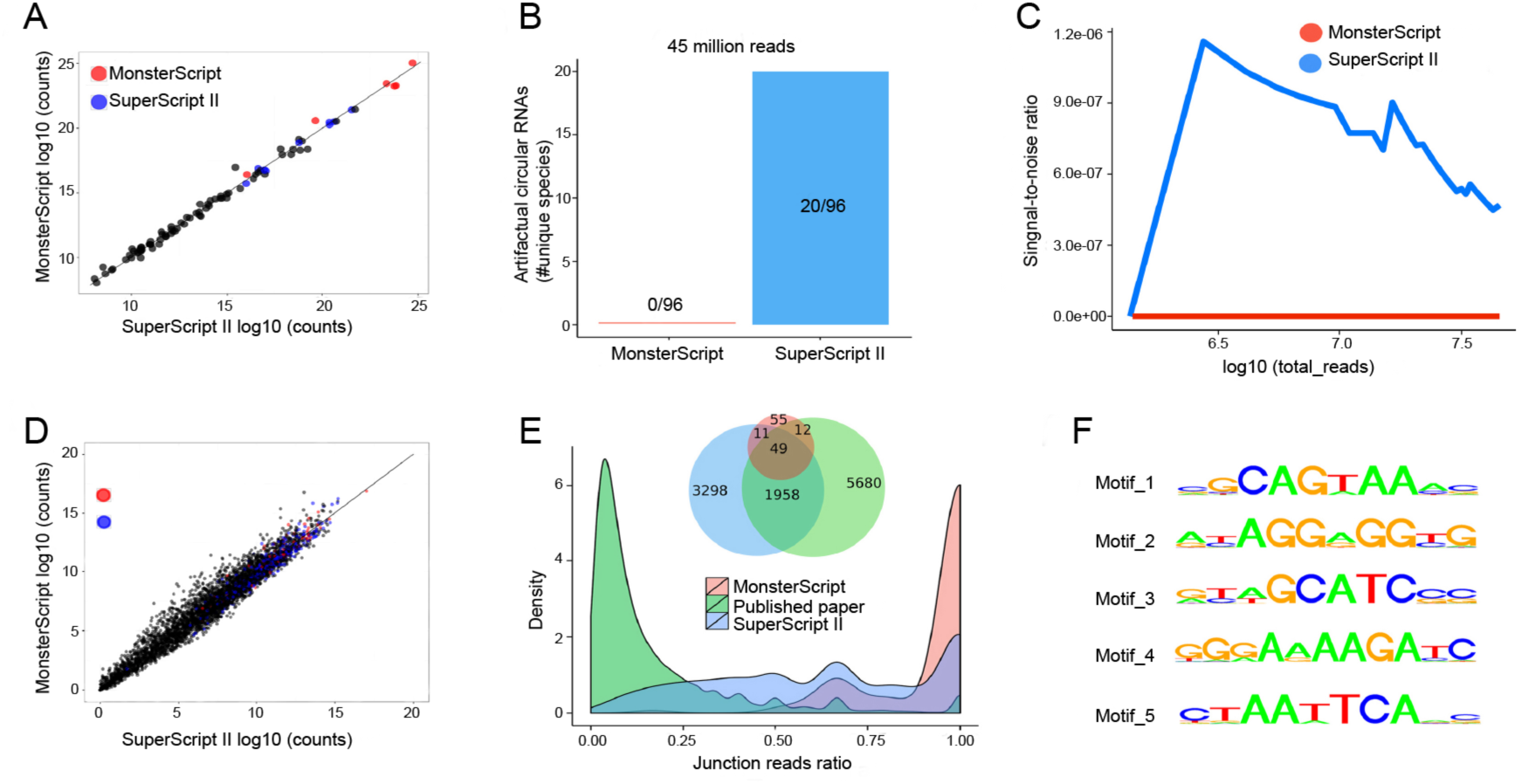
Deep sequencing analyses of control linear RNAs and RNAs from young (4-month-old) and old (2-year-old) brain samples. (**A**) Similar performance of RNA-Seq analyses on a library of 96 linear RNAs (ERCC RNA Spike-In Mix) using SuperScript II and MonsterScript. Data represent mean values of duplicated experiments. (**B**) Twenty artificial circular RNAs were identified from SuperScript II, but none from MonsterScript libraries of the ERCC RNA Spike-In Mix based on junction sequences. (**C**) Relationship between sequencing depth and noise-to-signal ratios of SuperScript II *vs.* MonsterScript-based library construction protocols for circular RNA identification. Sequencing data of the ERCC RNA Spike-In Mix were used for this analysis. (**D**) The scatter plot showing sequencing count distribution of all the genes detected in the duplicated libraries constructed using the two reverse transcriptases (MonsterScript and SuperScript II) using RNAs from brains samples of young (4-month-old) and old (2-year-old) mice. (**E**) Venn diagram showing the putative circular RNAs identified from mouse brain by a previous (8) and the present studies using SuperScript II- and MonsterScript-based library construction protocols. Junction reads ratios in mouse brain libraries from a previous and the present studies using SuperScript II and MonsterScript-based library construction protocols. (**F**) Potential motifs found in the junction sequences of artificial circular RNAs.

### Library construction using MonsterScript minimizes false identification of circular RNAs

To further evaluate the utility of the MonsterScript-based circular RNA identification protocol, we constructed mouse brain libraries using rRNA-depleted, RNase R-treated total RNAs followed by RT using MonsterScript and SuperScript II, respectively. Under similar sequencing depths (~50 million reads), both methods yielded similar read counts (**Fig. 2D**).

We then compared our circular RNA annotation results with those reported in an earlier report (8), which identified 7,638 putative circular RNAs from mouse brain using rRNA-depleted total RNAs and SuperScript II. Our analyses using age-matched, whole brain tissue of the same mouse strain identified 5,256 putative circular RNAs from SuperScript II libraries, among which 1,958 overlapped with those reported in the published paper (8). In sharp contrast, only 127 circular RNAs could be annotated from libraries constructed using MonsterScript (**Fig. 2E**), 61 of which were those reported in the published paper (8). These results suggest that the vast majority of the putative circular RNAs identified from SuperScript II-constructed libraries might be artifacts caused by template switching-induced junction sequences. Consistent with this notion, >90% of the putative circular RNAs identified in the previous report (8) displayed low junction reads ratios, which are, in general, suggestive of false positive (**Fig. 2E**). In contrast, the junction reads ratios in MonsterScript libraries were much higher (**Fig. 2E**). By analyzing the junction sequences of the artificial circular RNAs (i.e., those identified from SuperScript II-derived cDNA libraries of Spike-In linear RNAs) and the most likely true/biological circular RNAs (i.e., 72 circular RNAs shared between SuperScript II and MonsterScript brain libraries), we identified five significant motifs, all of which appeared to be GC-rich (**Fig. 2F**), a pattern well documented to be common in MMLV reverse transcriptases-induced template switching (9).

### High-resolution melting analyses can distinguish between circular RNAs from their linear forms

Although Northern blots have been regarded as a “gold standard” for circular RNAs validation (5), the method is less useful for low abundance or high-throughput analyses. As an alternative to Northern blots, we explored the utility of high-resolution melting (HRM) analyses to determine RNA circularity (**Fig. 3**). The HRM analyses have been previously employed to detect m6A nucleoside (10), and we modified the method such that a quencher probe quenches the fluorescence of a FAM probe when they anneal to the junction site of a circular RNA in a head-to-head orientation. During a gradual increase of temperature, the dark quencher oligo dissociates from the template at a specific temperature, thus allowing emission of fluorescence from the FAM probe (**Fig. 3A**). Since the quencher and the FAM probes anneal to the tail and head of the corresponding linear form, respectively, the fluorescence only weakly fluctuates without significant peaks during the melting process (**Fig. 3B**). Given that the heights of the fluorescence peaks correlate with the input of control circular RNAs (0.1 ng, 1ng and 10ng) in a linear fashion (**Fig. 3C**), the HRM analyses can also be used for quantitative analyses of circular RNAs based on the linear relationship between the height of the fluorescence peaks and the input of the control circular RNAs (**Fig. 3C**). Although this method can reliably distinguish RNA circles from their linear forms, specific probes need to be synthesized for individual circular RNAs, which are costly and may not be convenient for high throughput analyses. Nevertheless, the HRM analyses provide an alternative for distinguishing circular RNAs from their linear forms. This feature is important because it has been reported that many circular RNAs are co-expressed with their corresponding linear forms and the circular forms become much more enriched with aging (3, 8, 11), and there is no reliable method which can distinguish between circular and linear forms of the same sequences.

**Fig. 3.**
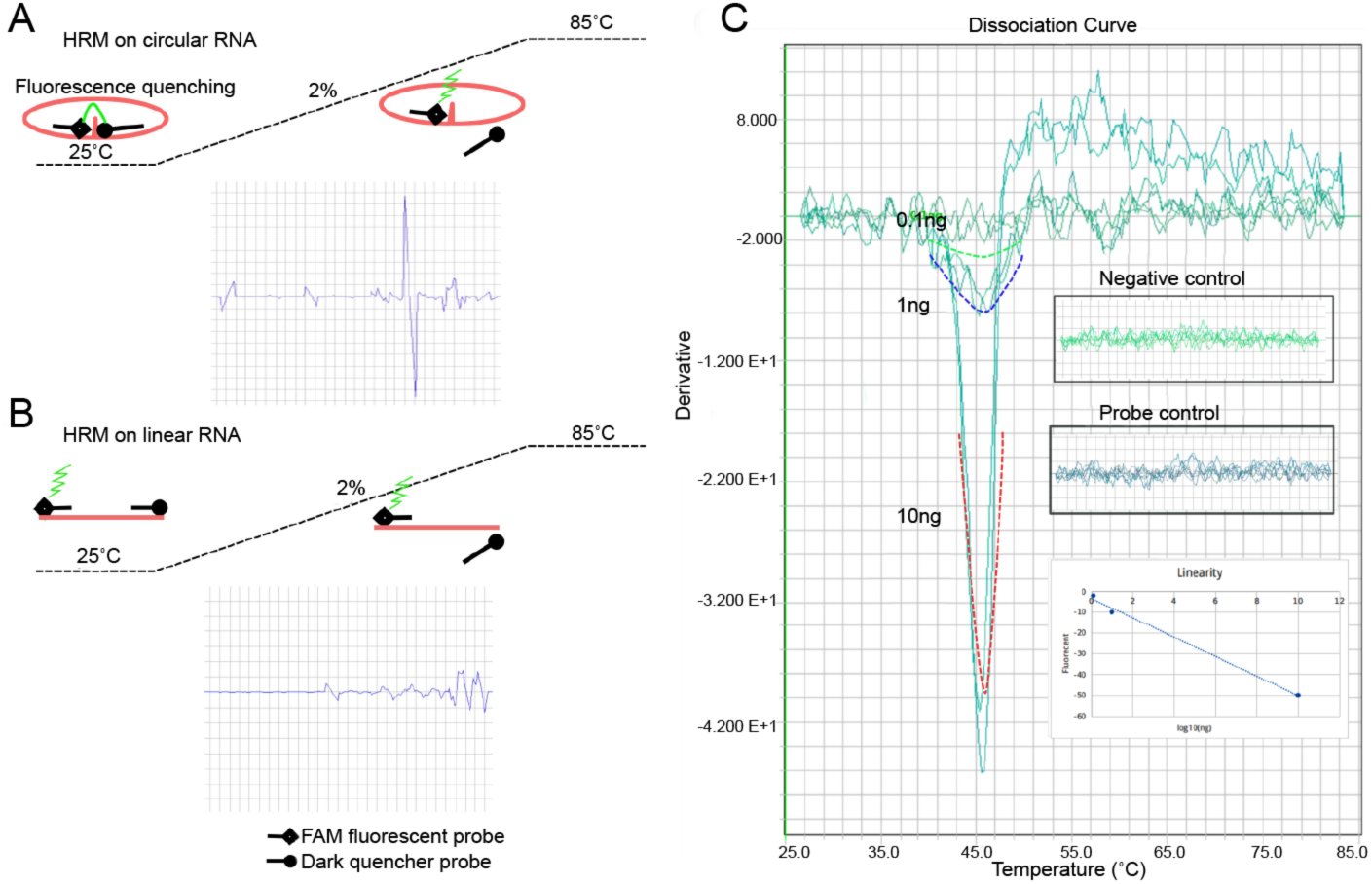
Detection and quantification of circular RNAs using high-resolution melting (HRM) analyses. (**A**) A dark quencher probe quenches the fluorescence of a FAM probe when they anneal to the junction site of a circular RNA in a head-to-head orientation. During a gradual increase of temperature, the dark quencher oligo dissociates from the template at a specific temperature, thus allowing emission of fluorescence from the FAM probe. Using our control circular RNA for HMR analyses, we observed a significant fluorescence peak at ~45°C. (**B**) Since the quencher and the FAM probes anneal to the tail and head of the corresponding linear form, respectively, the fluorescence only weakly fluctuates without significant peaks during the melting process. (**C**) HMR-based quantitative analyses of circular RNAs. Heights of the fluorescence peaks correlate with the input of the control circular RNAs (0.1 ng, 1ng and 10ng) in a linear fashion. No obvious peaks were observed in the negative (linear RNA) and probe-only (background) controls. Therefore, the HRM analyses can be used to quantify circular RNAs as well.

### Both MonsterScript junction PCR and high-resolution melting analyses can reliably determine RNA circularity

To test the robustness of the two new methods (i.e., MonsterScript junction PCR and HRM analyses), as compared to the traditional Northern blots, in determining the circularity of RNAs, we examined levels of a previously validated, brain-enriched circular RNA, mmu_circ_0011529(8), in old (2-year-old) and young (4-month-old) brain samples (**Fig. 4**). Junction PCR using total mouse brain cDNAs synthesized by SuperScript II detected mmu_circ_0011529 in both young and old brain samples with similar abundance, whereas this circular RNA was exclusively detectable in old mouse brain samples when MonsterScript-synthesized cDNAs were used for junction PCR (**Fig. 4A**). This result is consistent with that of previous Northern blots showing enrichment of this circular RNA in aged organs(3, 8). Northern blot analyses of mmu_circ_0011529 detected a band in old mouse brain, which was higher than that in young brain (**Fig. 4B**), suggesting that true circular RNA mmu_circ_0011529 and its linear form are expressed in old and young mouse brain, respectively. Further supporting this notion, HRM analyses detected circular RNA mmu_circ_0011529 in old brain, but not in young brain (**Fig. 4C**). Sanger sequencing of the PCR products from SuperScript and MonsterScript junction PCR (**Fig. 4A**) detected true mmu_circ_0011529 sequence in old brain cDNAs synthesized by either SuperScript or MonsterScript II, whereas an insertion was identified in SuperScript II junction PCR products from young brain, indicative of TS activity of SuperScript II (**Fig. 4D**). These data indicate that both MonsterScript junction PCR and HRM analyses can reliably distinguish circular RNAs from their linear forms.

**Fig. 4.**
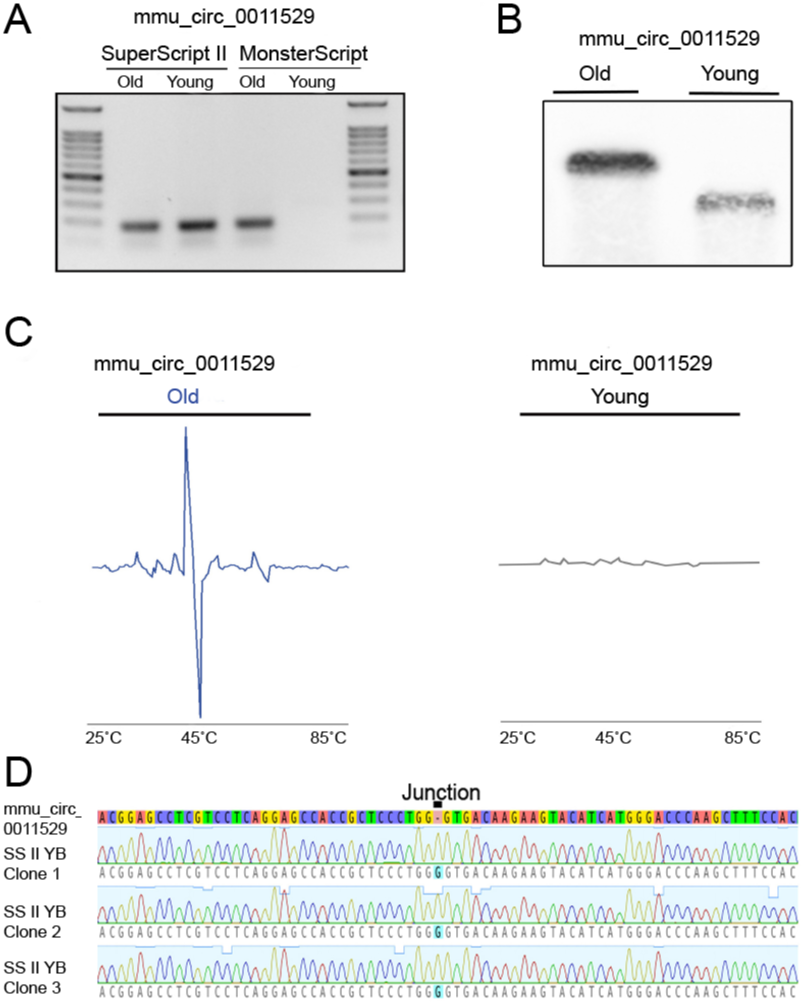
Validation of RNA circularity using Northern blot, MonsterScript junction PCR and HRM analyses. (**A**) Junction PCR using SuperScript II-synthesized cDNAs from RNase R-treated total RNA detected mmu_circ_0011529 in brain of both young (4-month-old) and old (2-year-old) mice with similar abundance, whereas this RNA was only detected in old, but not in young mouse brain when MonsterScript-synthesized cDNAs were used for junction PCR. (**B**) Northern blot analyses of mmu_circ_0011529 in old and young mouse brain samples using a probe from the junction site. Note that the band in the old brain sample was higher than that in the young brain sample, suggesting that the RNA detected in the old brain sample is the true circular RNA mmu_circ_0011529, whereas the one in the young brain sample is likely its linear form. (**C**) HRM analyses of mmu_circ_0011529 in old and young mouse brain samples. Circular RNA mmu_circ_0011529 was detected in old brain, but not in young brain. Note that the fluorescent peak results from disassociation of the quenching probe from the circular RNA at the junction site. (**D**) Sanger sequencing of the PCR products from SuperScript and MonsterScript junction PCR detected true mmu_circ_0011529 sequence in old brain cDNAs synthesized by either SuperScript or MonsterScript II, whereas an insertion was identified in SuperScript II junction PCR products from young brain, indicative of TS activity of SuperScript II.

We further analyzed >80 circular RNAs selected from the circBase based on their enrichment in the brain and testis of aged mice (3, 8, 11) *(SI Appendix, Fig. S3).* Interestingly, ~95% of these circular RNAs were detected in old testis and brain samples by MonsterScript junction PCR *(SI Appendix, Fig. S3),* suggesting that the aged testis and brain indeed express these RNA circles. However, although SuperScript junction PCR detected almost all of these RNAs in young testis and brain samples, MonsterScript junction PCR could only confirm a small fraction (<30%) *(SI Appendix, Fig. S3),* supporting the notion that these RNAs are expressed as linear RNAs in young mice, but become RNA circles with aging. Taken together, our data suggest that in addition to Northern blots, both MonsterScript junction PCR and HRM can be alternative methods for determining circularity of large RNAs.

In summary, we experimentally demonstrated that the commonly used reverse transcriptases can cause template switching, leading to false positive identification of junction sequences and consequently misclassification of linear RNAs as RNA circles. The problem can be avoided, at least partially, by using MonsterScript for library construction and circular RNA discovery, and MonsterScript junction RT-PCR in conjunction with HRM for circular RNA validation.

## Materials and Methods

### Animals

All mice used in this study were on the C57BL/6 J background, and housed under specific pathogen-free conditions in a temperature- and humidity-controlled animal facility at the University of Nevada, Reno. Animal use protocol was approved by Institutional Animal Care and Use Committee (IACUC) of the University of Nevada, Reno (Protocol number 00494), and are in accordance with the “Guide for the Care and Use of Experimental Animals” established by National Institutes of Health (NIH) (1996, revised 2011).

### Preparation of RNA

Total RNA was isolated using the mirVana total RNA extraction kit (Cat#: AM1560, Ambion) following the manufacturer’s protocol. The HiScribe T7 High-Yield RNA Synthesis Kit (Cat#: E2040S, NEB) was used for *in vitro* transcription to synthesize the control linear RNAs with the FLuc control plasmid as the template. The control linear RNAs were circulated using T4 RNA ligase 1 (Cat#M0204S, NEB) to generate circular RNAs. Sequences of the control plasmid, linear and circular RNAs can be found in *SI Appendix, File S1.*

### RT-PCR analyses of linear and circular RNAs

Cellular RNAs or synthetic control linear or circular RNAs were reverse transcribed using MonsterScript (Cat# MS041050, Epicentre) and SuperScript II (Cat# 18064014, Thermo Fisher) in a reaction containing the following reagents: 4μl 5xBuffer (supplementing 10mM DTT for SuperScript II), 1μl dNTP, 1μl reverse transcriptase, 1μl control RNA, 1μl Smarter IIA oligo (10μM), 1μl gene-specific primer (10uM), 11μl water. The reverse transcription was performed in 25°C 10min, 42°C 30 min, 65°C 30min, 85°C 5min. PCR was performed using GoTaq DNA polymerase (Cat# M3001, Promega) and convergent/divergent primers at the annealing temperature of 55°C. The PCR products were visualized with agarose gel electrophoresis. Primer sequences can be found in *SI Appendix, Table S1.* The DNA ladder is 100bp DNA Ladder H3 RTU (GeneDireX).

### RNase R treatment

Control linear and circular RNAs (100ng) were incubated with or without 20U of RNase R (Cat#RNR07250, Epicentre) in the reaction buffer for 1h at 37°C. The treated RNA samples were purified using the RNeasy Mini Kit (Cat#74104, Qiagen) and quantified. Equal amounts of RNA were then subjected to reverse transcription using the SuperScript II or MonsterScript and the random hexamer followed by PCR.

### Reverse transcriptase sensitivity assay

For measuring the relative efficiency of the two reverse transcriptases (SuperScript II and MonsterScript), the control linear RNA or RNase R-treated brain total RNAs were diluted to generate serial 5/10-fold dilutions, ranging from 5ng to 1pg. Reverse transcription was performed using MonsterScript and SuperScript II followed by real-time PCR on an ABI 7900HT qPCR system using convergent primers. The Ct values were converted to log 10 values for generating the linear regression plots.

### RNA-seq

For RNA-seq analyses, SuperScript II libraries were constructed using the KAPA Stranded RNA-Seq Kit (Cat#KK8483, Kapabiosystems) after ribosome depletion with RiboErase (Cat#KK8483, Kapabiosystems). To construct MonsterScript libraries for RNA-seq, SuperScript II included in the Kit was substituted with MonsterScript. The linear spike-in control was purchased from Thermo Fisher (Cat#4456740). All the samples were run in the same lane for deep sequencing on an Illumina NextSeq sequencer with PE75 in the Nevada Genomic Center.

### Linear RNA annotation

The sequence reads of the control linear RNAs (i.e., spike-in RNAs) was mapped to the reference sequences by Tophat2(12). RNA-seq reads of mouse tissues were mapped to the UCSC mm9 genome by Tophat2(12). The mapped bam files were counted though Subread/featureCounts(13). The counts of each gene were imported into Deseq2(14), followed by normalization through regularized logarithm (rlog). We used the ggplot2 packages(15) to generate plots.

### Circular RNA annotation

Sequencing reads of the spike-in control were mapped to spike-in reference sequences using bwa (bwa mem -T 19)(16). RNA-seq reads of mouse brain were mapped to UCSC mm9 genome by bwa (bwa mem -T 19(16). CIRI2(17) was used for circular RNA annotation (CIRI_v2.0.1.pl --no_strigency -I bwa.sam -O ciriout -F bwaindex/genome.fa).

### Motif analysis

The supported circular RNA reads were extracted from raw data. The junction reads with lowest junction reads ratio (<0.01) were likely the false positive ones in the spike-in RNA-Seq analyses. The junction reads with highest junction reads ratio (>0.7) were mostly the true positive reads (MonsterScript RNA-Seq on the aged mouse brain). We analyze the template-switching motifs using false-positive reads against the control, true positive reads. The motif analyses were performed using *motifRG*(18). Five significant motifs were identified and all appeared to be GC-rich, a pattern that has been documented to be common in MMLV reverse transcriptase-induced template switching(9).

### Northern blot

The Northern blot experiments were performed using NorthernMax Kit (Ambion) and detected using Biotin Chromogenic Detection Kit (Thermo Scientific), following the manufactures’ instructions. For the sensitivity experiment, biotin-labeled DNA probes spanning the whole control RNA was generated using Biotin DecaLabel DNA Labeling Kit (Thermo Scientific), following the manufacture’s recommended protocol. The FLuc linear DNA plasmid was used as template for probe generation. Heat denatured 1.8kb control FLuc RNA was used. For Northern blot using single junction probes, single biotin-labeled oligo probes were ordered from IDT and used to detect the linear control RNA. For Northern blot detection of circular RNA mmu_circ_0011529, a mixture of 8 biotin-labeled probes were use. 2µg of mouse total brain RNA were loaded on each lane for each sample. For probe sequences please see *SI Appendix, Table S2.*

### High-resolution melting (HRM) analysis

The probe mix contains 10µM FAM probe and 20µM quencher probe. RNA was mixed with 2µl of probe mix, 10µl of 10X hybridization buffer (1 M NaCl, 0.1 M Tris-HCl), and RNase-free water to a final concentration of 100µl. The samples were analyzed in 7900HT Fast Real-Time PCR System (Applied Biosystems) and incubated as follows: 75˚C for 5min, cool down to 25˚C for 15s, ramp heating to 85˚C at 2%. Data collection was set during the ramp heating and at 85˚C.

## Acknowledgements

This work was supported by grants from the NIH (HD071736 and HD085506 to WY) and the Templeton Foundation (PID: 50183 to WY). RNA-Seq and bioinformatics were conducted in the Single Cell Genomics Core of University of Nevada, Reno School of Medicine, which was supported, in part, by a COBRE grant from the NIH (1P30GM110767).

## Author Contributions

C.T. and W.Y. designed the research. C.T., T.Y., Y.Z. and H.Z. performed the experiments. All participated in data analyses. C.T. and W.Y. wrote the manuscript. All reviewed the manuscript.

## References

1. Capel B, et al. (1993) Circular transcripts of the testis-determining gene Sry in adult mouse testis. Cell 73(5): 1019-1030.

2. Jeck WR & Sharpless NE (2014) Detecting and characterizing circular RNAs. Nat Biotechnol 32(5):453-461.

3. Memczak S, et al. (2013) Circular RNAs are a large class of animal RNAs with regulatory potency. Nature 495(7441):333-338.

4. Hansen TB, Veno MT, Damgaard CK, & Kjems J (2016) Comparison of circular RNA prediction tools. Nucleic Acids Res 44(6):e58.

5. Szabo L & Salzman J (2016) Detecting circular RNAs: bioinformatic and experimental challenges. Nat Rev Genet 17(11):679-692.

6. Panda AC, et al. (2017) High-purity circular RNA isolation method (RPAD) reveals vast collection of intronic circRNAs. Nucleic Acids Res.

7. AbouHaidar MG & Ivanov IG (1999) Non-enzymatic RNA hydrolysis promoted by the combined catalytic activity of buffers and magnesium ions. Z Naturforsch C 54(7–8):542-548.

8. You X, et al. (2015) Neural circular RNAs are derived from synaptic genes and regulated by development and plasticity. Nat Neurosci 18(4):603-610.

9. Zajac P, Islam S, Hochgerner H, Lonnerberg P, & Linnarsson S (2013) Base preferences in non-templated nucleotide incorporation by MMLV-derived reverse transcriptases. PLoS One 8(12):e85270.

10. Golovina AY, et al. (2014) Method for site-specific detection of m6A nucleoside presence in RNA based on high-resolution melting (HRM) analysis. Nucleic Acids Res 42(4):e27.

11. Glazar P, Papavasileiou P, & Rajewsky N (2014) circBase: a database for circular RNAs. RNA 20(11):1666-1670.

12. Trapnell C, Pachter L, & Salzberg SL (2009) TopHat: discovering splice junctions with RNA-Seq. Bioinformatics 25(9): 1105-1111.

13. Liao Y, Smyth GK, & Shi W (2014) featureCounts: an efficient general purpose program for assigning sequence reads to genomic features. Bioinformatics 30(7):923-930.

14. Love MI, Huber W, & Anders S (2014) Moderated estimation of fold change and dispersion for RNA-seq data with DESeq2. Genome Biol 15(12):550.

15. Wickham H (2009) ggplot2: Elegant Graphics for Data Analysis. (Springer-Verlag, New York).

16. Li H & Durbin R (2009) Fast and accurate short read alignment with Burrows-Wheeler transform. Bioinformatics 25(14):1754-1760.

17. Gao Y, Wang J, & Zhao F (2015) CIRI: an efficient and unbiased algorithm for de novo circular RNA identification. Genome Biol 16:4.

18. Yao Z, et al. (2014) Discriminative motif analysis of high-throughput dataset. Bioinformatics 30(6):775-783.

